# Yeast-Expressed SARS-CoV Recombinant Receptor-Binding Domain (RBD219-N1) Formulated with Aluminum Hydroxide Induces Protective Immunity and Reduces Immune Enhancement

**DOI:** 10.1101/2020.05.15.098079

**Authors:** Wen-Hsiang Chen, Xinrong Tao, Anurodh Agrawal, Abdullah Algaissi, Bi-Hung Peng, Jeroen Pollet, Ulrich Strych, Maria Elena Bottazzi, Peter J. Hotez, Sara Lustigman, Lanying Du, Shibo Jiang, Chien-Te K. Tseng

## Abstract

We developed a severe acute respiratory syndrome (SARS) subunit recombinant protein vaccine candidate based on a high-yielding, yeast-engineered, receptor-binding domain (RBD219-N1) of the SARS beta-coronavirus (SARS-CoV) spike (S) protein. When formulated with Alhydrogel®, RBD219-N1 induced high-level neutralizing antibodies against both pseudotyped virus and a clinical (mouse-adapted) isolate of SARS-CoV. Here, we report that mice immunized with RBD219-N1/Alhydrogel® were fully protected from lethal SARS-CoV challenge (0% mortality), compared to ∼ 30% mortality in mice when immunized with the SARS S protein formulated with Alhydrogel®, and 100% mortality in negative controls. An RBD219-N1 formulation Alhydrogel® was also superior to the S protein, unadjuvanted RBD, and AddaVax (MF59-like adjuvant)-formulated RBD in inducing specific antibodies and preventing cellular infiltrates in the lungs upon SARS-CoV challenge. Specifically, a formulation with a 1:25 ratio of RBD219-N1 to Alhydrogel® provided high neutralizing antibody titers, 100% protection with non-detectable viral loads with minimal or no eosinophilic pulmonary infiltrates. As a result, this vaccine formulation is under consideration for further development against SARS-CoV and potentially other emerging and re-emerging beta-CoVs such as SARS-CoV-2.

## INTRODUCTION

Coronaviruses (CoV) are the enveloped viruses with approximately 30 kb single-strand RNA genomes. CoVs belong to the family Coronaviridae and have been found in various mammals, including bats, pangolins, and civets. Previously, they were known to only cause mild diseases to humans until the pandemic of severe acute respiratory syndrome (SARS) occurred between 2002 and 2003 [1-3]. Ever since SARS, nearly every decade, a new major coronavirus outbreak occurred: The Middle East respiratory syndrome caused by MERS-CoV first emerged in 2012 and still is circulating in camels [4]; the current COVID-19 pandemic caused by SARS-CoV-2 was first discovered in December 2019, and have currently infected more than 10 million worldwide.

The disease caused by SARS coronavirus (SARS-CoV) led to almost 800 deaths and more than 8,000 infections, leading to an overall fatality rate of approximately 10 percent. Alarmingly, the fatality rate among older adults exceeded 50 percent [5]. In preparation for future outbreaks and accidental and/or intentional releases of SARS-CoV, intensive efforts have been made to develop vaccines against SARS-CoV.

For the past two decades, several antigens have been identified and developed as SARS-CoV vaccine candidates. Initially, whole inactivated virus (WIV) or modified vaccinia virus Ankara expressing SARS vaccines were developed [5], however, eosinophilic immunopathology was observed in mice and non-human primates immunized with these viral-vectored vaccines [6-11]. Even though historically there have been reports that alum adjuvanted vaccines could induce enhancement, such as in the 1960s with the RSV vaccine or even with WIV and S proteins,[12] it was shown later that alum-adjuvanted WIV elicited less immunopathology than WIV alone [12]. suggesting that alum might reduce immune enhancement, a process possibly linked to mixed Th1, Th17, and Th2 responses [11, 13]. Additional evidence emerged that the virus N protein had a key but not exclusive role in immune enhancement [7, 11]. Based on these studies, the recombinant S protein of SARS-CoV was used as a vaccine candidate [5], but the full-length S-protein also induced immunopathology, with epitopes outside of the receptor-binding domain (RBD) of the S protein implicated in eliciting this phenomenon [14, 15]. Therefore, the RBD of the S protein was selected as a substitute for the full-length S protein [16-21].

Recombinant RBD formulated with Sigma’s adjuvant system® (consisting of Monophosphoryl-lipid A/Trehalose dicorynomycolate adjuvant, a skewed Th1/Th2 adjuvant) or with Freund’s adjuvant (Freund’s complete in prime and Freund’s incomplete in boost; a Th1/Th2 balanced adjuvant scheme) was shown to elicit neutralizing antibodies and highly protective immunity in the vaccinated animals, while eliminating or significantly reducing eosinophilic immunopathology [18, 20-23]. In our previous studies, we have expressed wild-type RBD193 (residues 318–510) /RBD219 (residues 318–536) in yeast, however, because of the three N glycosylation sites on these two wild-type constructs, we further generated the deglycosylated forms as follows: N1: 1^st^ glycosylation site deleted; N2: 1^st^glycosylation site deleted and 2^nd^ glycosylation site mutated; and N3: 1^st^ glycosylation site deleted, 2^nd^and 3^rd^glycosylation sites mutated. We developed a production process for several of these tag-free yeast-expressed recombinant RBD constructs [24]. Such studies down-selected several candidates and ultimately identifying one, RBD219-N1 (residues 319-536), as a promising vaccine candidate, due to its ability to induce in immunized mice a stronger anti-RBD-specific antibody response and neutralizing antibodies when adjuvanted with aluminum hydroxide (Alhydrogel®). The protein production process [25] was transferred to the Pilot Bioproduction Facility (PBF) at Walter Reed Army Institute of Research (WRAIR), and the clinical grade RBD219-N1 (drug substance) was manufactured under current Good Manufacturing Practices (cGMP) and is suitable for further Phase I clinical trials.

In this work, the RBD219-N1 formulated with Alhydrogel® resulted in significantly increased antigen-specific IgG titers and neutralizing antibody responses when compared to other RBD constructs. After challenge with SARS-CoV, 100% of mice immunized with RBD219-N1 survived, while only 89% of mice immunized with other RBD constructs and less than 70% of the mice immunized with SARS-CoV spike protein survived and none survived in the control group. The aluminum formulated RBD minimized immune enhancement compared with other adjuvants formulations with either the RBD or full-length S protein. Finally, an Alhydrogel® dose-ranging study further indicated that by formulating RBD219-N1 with Alhydrogel® at the ratio of 1:25, higher IgG titers could be elicited with no detectable viral load upon challenge.

## MATERIALS AND METHODS

### Generation of recombinant yeast-expressed RBD of SARS-CoV

The yeast-expressed SARS-CoV RBD193-N1, wt-RBD219, and RBD219-N1 were expressed and purified as previously described [24, 25]. Briefly, 1 mL of *P. pastoris* X33 seed stock expressing RBD193-N1, wt RBD219, and RBD219-N1 was inoculated into 500 ml BMG (buffered minimal glycerol) medium and the culture was incubated overnight at 30°C with constant shaking at 250 rpm until an OD600 of ∼10. Approximately 250 ml of overnight culture were inoculated into 5 L sterile Basal Salt Media or Low Salt medium [24]. Fermentation was maintained at 30°C, pH 5.0 and 30% of dissolved oxygen concentration until the exhaustion of glycerol, and the pH and the temperature were then ramped to 6.5 and 25°C, respectively, over an hour followed by continuous feeding of methanol at 11 ml/L/hr for ∼70 hours. The fermentation supernatant (FS) was harvested for further purification. To purify RBD193-N1, wt-RBD219, and RBD219-N1, ammonium sulfate was added to the FS until the molarity reached 2 M. The FS containing 2 M ammonium sulfate was purified by hydrophobic interaction chromatography using Butyl Sepharose HP resin followed by size exclusion chromatography using Superdex 75 resin [24, 25].

### Reagents

Alhydrogel® (aluminum oxyhydroxide; Catalog # 250-843261 EP) was purchased from Brenntag (Ballerup, Denmark), AddaVax (MF59-like adjuvant; squalene oil-in-water emulsion; Catalog # vac-adx-10) was purchased from Invivogen (San Diego, CA, USA). The SARS S protein vaccine, produced in the baculovirus/insect cell expression platform and pre-formulated with aluminum (Reagent # 50-09014, 50-09015, 50-09016), was obtained directly from NIH via BEI Resources, NIAID, NIH (Manassas, VA, USA).

### Binding Study

One ml of TBS containing 18 to 180 µg RBD219-N1 and 400 µg Alhydrogel® were prepared to study the binding of RBD219-N1 to Alhydrogel® at different ratios (from 1:2 to 1:22). The prepared RBD219-N1/ Alhydrogel®slurry was mixed for one hour to ensure the binding of RBD219-N1 to Alhydrogel® reached an equilibrium state. The slurry was then centrifuged at 13,000 x g for 5 minutes, and the supernatant was collected while the Alhydrogel®pellet was resuspended with an equal volume of removed supernatant. The RBD219-N1 protein content in the supernatant fraction and the pellet fraction were then measured using a micro BCA assay (ThermoFisher, Waltham, MS, USA). Similarly, the presence of RBD219-N1 in the pellet and supernatant fractions was also evaluated using SDS-PAGE. Briefly, after the slurry was centrifuged and separated into pellet and supernatant fractions, the Alhydrogel® pellet was further resuspended with desorption buffer (100 mM sodium citrate, 92 mM dibasic sodium phosphate at pH 8.9) and mixed for 1 hour. The desorbed RBD was then separated from Alhydrogel® by centrifugation at 13,000 x g for 5 minutes. Ten microliters of desorbed RBD from the pellet fraction and free RBD in the supernatant fraction were loaded on 4-20% Tris-glycine gels and stained by Coomassie blue.

### Animals

Six- to eight-week-old, female Balb/c mice were purchased from Charles River (Wilmington, MA, USA), and housed in an approved biosafety level 3 animal facility at the University of Texas Medical Branch (UTMB) at Galveston, Texas. All of the experiments were performed according to National Institutes of Health and United States Department of Agriculture guidelines using experimental protocols approved by the Office of Research Project Protections, Institutional Animal Care and Use Committee (IACUC) at UTMB.

### Study Design

Three sets of pre-clinical studies were conducted to evaluate: (1) different yeast-expressed recombinant antigens, including RBD193-N1, wt-RBD219, and RBD219-N1), and spike protein vaccine as a control; (2) two of the most common adjuvants used in licensed vaccines, Alhydrogel®® and AddaVax (MF59-like adjuvant); and (3) two Alhydrogel® doses (1:8 and 1:25 ratios) to formulate RBD219-N1, and spike protein vaccine as a control. Mice were immunized either subcutaneously (s.c.) or intramuscularly (i.m.) followed by two boosters at 21-day intervals. The same route, number, and frequency of immunization were followed among all the groups within the same study. Mouse-adapted SARS-CoV (MA15 strain) was used in these studies to challenge the mice. This virus was generated by serially passaged in the respiratory tract of young BALB/c mice, resulting in minimal mutations in only 6 amino acids, and has been reported to show dose-dependent weight loss and mortality as well as associated pulmonary histopathology in BALB/c mice [26], and thus is widely used in a mouse model to evaluate SARS-CoV vaccine and therapeutics. The detailed study designs are described as the following:

#### Antigens screening

Mice (15 per group) were immunized s.c. with 100 µL yeast-expressed recombinant RBD proteins (RBD193-N1, wt-RBD219, and RBD219-N1) formulated with 10 mg/mL Alhydrogel® on days 0 (20 µg RBD), 21 (10 µg RBD), and 42 (10 µg RBD). TBS/Alhydrogel® buffer and 9 µg of alum pre-formulated SARS S protein were used as the negative and positive controls, respectively. Sera from 5 mice were collected on day 50 to evaluate the pre-challenge neutralizing antibody titers. All mice were then challenged intranasally (i.n.) with 10x LD50 SARS-CoV MA15 virus (∼5.6 logs TCID50) on day 52. Three mice in each group were further sacrificed On days 55 and 56 to determine the viral loads. The remaining 9 mice in each group were used to monitor clinical disease (weight loss) and mortality daily for up to three weeks.

#### Adjuvant screening

Mice (4 per group) were immunized intramuscularly (i.m.) with 100 µL RBD219-N1 formulated with two adjuvants on days 0 (20 µg RBD), 21 (10 µg RBD), and 42 (10 µg RBD). A total of three groups were tested, including RBD219-N1 with 500 µg Alhydrogel® (sometimes referred to in the manuscript as “alum”) in group 1, RBD219-N1 in 50% (v/v) MF59-like adjuvant in group 2. Mice immunized with RBD alone in group 3 were used as a negative control. On day 52, sera were collected to evaluate IgG and neutralizing antibody titers. On day 63, mice were i.n. challenged with SARS-CoV (2x LD50 (∼10^5^ TCID50) SARS-CoV (MA-15), and finally, on day 66 and day 69, or day 3 and day 6 post-challenge, lungs from 2 mice were collected for histopathology and viral titration.

#### Alhydrogel® dose range screening

Mice (6 per group) were i.m. immunized with Alhydrogel® formulated RBD219-N1 on days 0 (10 or 20 µg RBD), 21 (10 µg RBD), and 42 (10 µg RBD). In group 1, a formulation of 0.2 mg/mL RBD in 5 mg/mL Alhydrogel® (2.5 mg/mL aluminum) with the dose of 20 µg/10 µg/10 µg of RBD for prime/1st boost/2nd boost was used for immunization, respectively. In group 2, the formulation0.2 mg/mL RBD in 1.6 mg/mL Alhydrogel® (consisted of 0.8 mg/mL aluminum) was tested, more specifically, a dosing regimen of 20 µg/10 µg/10 µg RBD for prime/1st boost/2nd boost was used while in group 3, 10 µg RBD were used for all immunizations. Alhydrogel® alone and pre-formulated SARS-S protein (3 µg) were used in groups 4 and 5, respectively, as negative and positive controls. On day 52, sera were collected to evaluate IgG and neutralizing antibody titers. On day 63, mice were i.n. challenged with SARS-CoV (2x LD50 (∼10^5^ TCID50) SARS-CoV (MA-15)), and finally, on day 66 (day 3 post-challenge) and day 70 (day 6 post-challenge), lungs from 3 mice were collected for histopathology and viral titration.

### ELISA

RBD-specific IgG titers of polyclonal sera from the immunized mice were measured by ELISA, as previously described [18, 19, 24]. Briefly, 96-well ELISA plates were pre-coated with yeast-expressed RBD protein (1 µg/ml) overnight at 4 °C. After blocking and then incubating with serially diluted mouse sera, bound IgG antibody was detected using HRP-conjugated anti-mouse IgG (1:2000), followed by the same protocol as described [18, 19, 24].

### Titration of SARS CoV-specific neutralizing antibodies

Mice were anesthetized with isoflurane and then bled from the retro-orbital sinus plexus. After heat inactivation at 56 °C for 30 min, sera were stored at 4 °C. The standard live virus-based microneutralization (MN) assay was used as previously described [12, 27]. Briefly, serially two-fold and duplicate dilutions of individual immune sera were prepared in 96-well microtiter plates with a final volume of 60 μl per well before adding 120 infectious SARS-CoV (MA-15) particle in 60 μl to individual wells. The plates were mixed well and cultured at room temperature for 1 h before transferring 100 μl of the immune serum-virus mixtures into designated wells of confluent Vero E6 cells grown in 96-well microtiter plates. Vero E6 cells cultured with medium with or without virus were included as positive and negative controls, respectively. After cultivation at 37°C for 3 days, individual wells were observed under the microcopy for the status of virus-induced formation of cytopathic effect. The efficacy of individual sera was calculated and expressed as the lowest concentration capable of completely preventing virus-induced cytopathic effect in 100% of the wells.

### Collection of lungs, histology, immunohistochemistry, and virus titration

After the SARS-CoV challenge, mice were euthanized on different days depending on the study, and their lungs were removed. Lung lobes were placed in 10% neutral buffered formalin for histological examination using either hematoxylin and eosin (for cellular infiltrates) or immunohistochemistry (IHC), specific for eosinophils, as described previously[12, 27]. For virus quantitation, the remaining tissue specimen was processed as previously described [12, 27]. Evaluations for histopathology were done by an experimental human pathologist masked as to the vaccine/dosage of each specimen source; assessment of the extent of pathologic damage and the eosinophilic component of the inflammatory infiltrates was then provided.

### Statistical analysis

Neutralizing antibody titers, weight loss, lung virus titers, IgG titers, histopathologic score, and eosinophilic infiltration scores were averaged for each group of mice. T-tests were routinely used to evaluate the statistical variation between two groups.

## RESULTS

### Vaccine integrity evaluation through binding and point-of-injection studies

As a vaccine, recombinant protein antigens alone often do not induce a sufficient immune response, necessitating their evaluation as proteins formulated with adjuvants. When Alhydrogel® is used as an adjuvant, the antigen is typically fully adsorbed to the alum salt to maximize potency. The binding efficiency of RBD219-N1 to Alhydrogel® was performed to evaluate the minimum RBD219-N1 to Alhydrogel® ratio required to ensure complete protein binding. By measuring the protein content in the supernatant and the pellet fraction after adsorption and centrifugation, the percentage of RBD bound onto Alhydrogel® at different RBD219-N1 to Alhydrogel® ratio was determined (Figure 1A). An Alhydrogel® to RBD ratio greater than 7.4 resulted in complete adsorption. SDS-PAGE analysis of a formulation of 0.2 mg/mL RBD with 1.6 mg/mL Alhydrogel® (1:8 ratio) further confirmed that no protein remained in the supernatant fraction after adsorption (Figure 1B).

**Figure 1.**
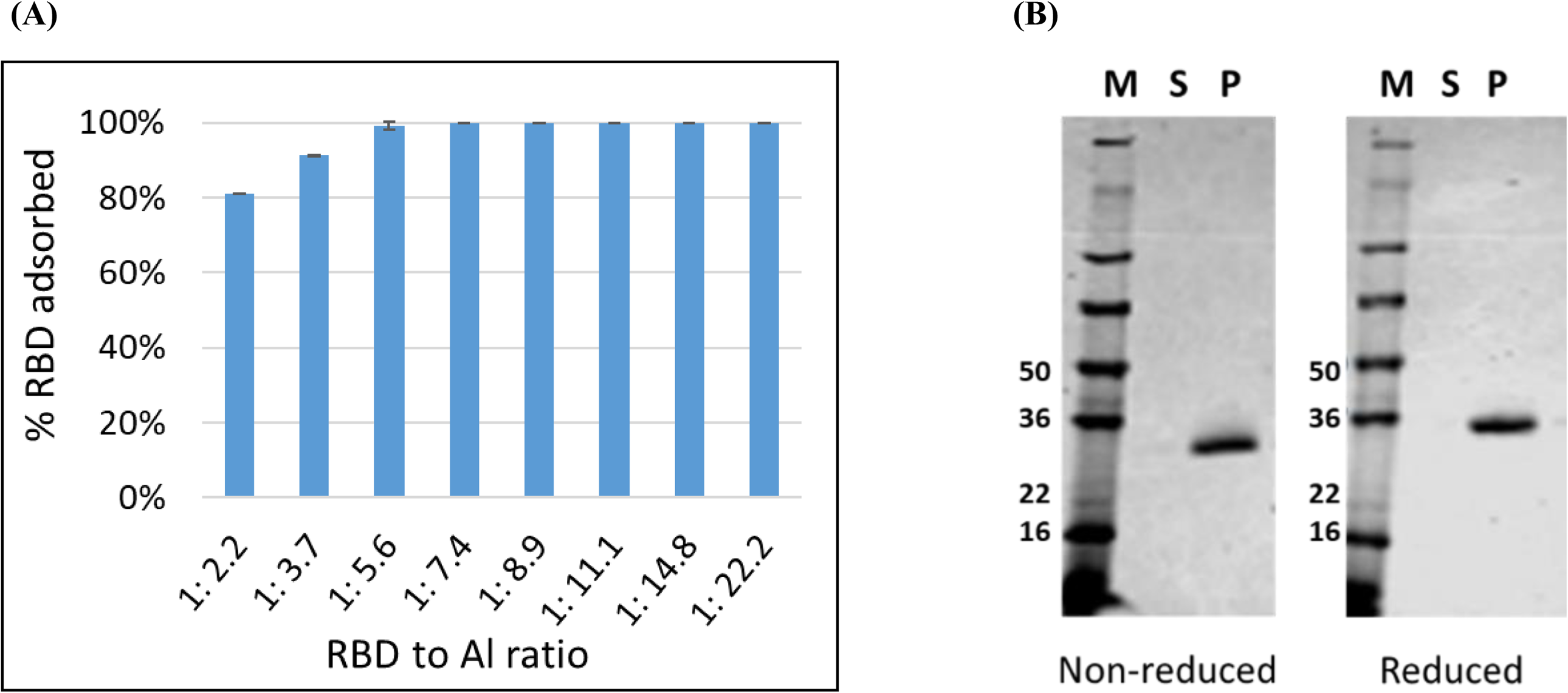
RBD to Alhydrogel® binding analysis. (A) A micro BCA assay was used to quantify the % RBD adsorbed on Alhydrogel® at different RBD to Alhydrogel® (Al) ratios; (B) SDS-PAGE analysis: the supernatant and pellet fractions for a formulation of 0.2 mg/mL RBD219-N1 with 1.6 mg/mL Alhydrogel (1:8 ratio) were analyzed. 10 µL of the desorbed RBD from the pellet fraction (P) and 10 µL of supernatant (S) were loaded on the gel and stained with Coomassie Blue. M: protein marker.

### Antigen screening

In this pilot study, we compared safety, immunogenicity, and efficacy of the various yeast-expressed RBD-and S vaccines [12] using a lethal mouse model of SARS--CoV infection. Figure 2a shows that mice immunized with RBD-219N1 had the highest titer of neutralizing antibodies compared to mice immunized with alum-adjuvanted RBD193-N1, wt-RBD219, and the S protein vaccines. Importantly, endpoint evaluation for mortality has shown that a 100% survival rate was found for mice immunized with alum-adjuvanted RBD219-N1, while mice immunized with alum-adjuvanted wt-RBD219 and RBD193-N1 vaccines showed 88% survival and those immunized with alum-adjuvanted S protein showed only 67% survival. All mice in the TBS control group died within 6 days post-challenge (Figure 2B). Furthermore, mice immunized with RBD219-N1, similar to wt-RBD219 and RBD193-N1, consistently showed less than 10% weight loss throughout the study period, while mice immunized with other alum-formulated vaccines including the S protein showed a maximum of 15-20% weight loss (Figure 2C). This was accompanied by more than a 3-log reduction of infectious viral loads within the lungs when compared to mice vaccinated with TBS/Alhydrogel® (Figures 2D and 2E). None of the mice given TBS/Alhydrogel® produced detectable neutralizing antibodies, whereas their geometric means of lung virus titers were 10^9.9^ and 10^8.9^ TCID_50_/g on days 1 and 2 post-challenge, respectively. With these results, RBD219-N1 was chosen for further development.

**Figure 2.**
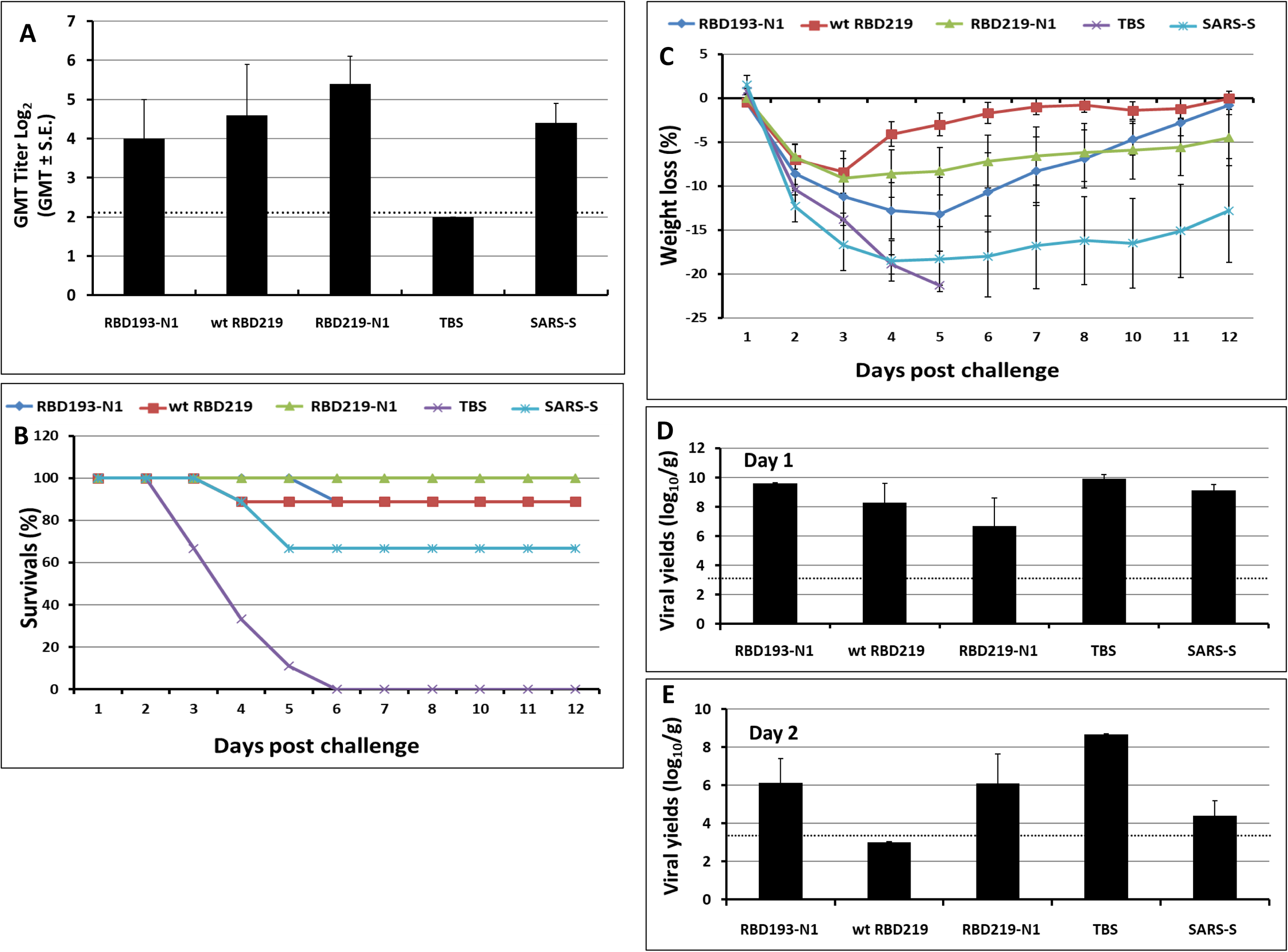
Vaccination-induced protection against lethal MA-15 infection at different stages. Post immunization: (A) Neutralizing antibody titers after immunization. Post challenge: (B) daily survival rates, (C) daily weight loss, and the viral loads in the lung on day 1 (D) and day 2 (E).Groups of mice (N=15 per group) were immunized 3 times with yeast expressed RBDs (20, 10, and 10 µg respectively) or 9 µg of S protein for each immunization at 3-week intervals. Mice given TBS/alum were included as controls. The titers of neutralization antibodies were determined on day 50. All vaccinated mice were challenged with 5.6 logs (∼ 10X LD_50_) TCID_50_/60 µL of MA-15 intranasally (IN). Three challenged mice in each group were euthanized on days 1 and 2 post-challenge, respectively. The remaining mice in each group (N=9) were monitored daily for clinical manifestations (e.g., weight loss), and mortality.

### Adjuvant screening

It is known that alum (generally a Th2 adjuvant) and MF59 (a Th1/Th2 balanced adjuvant in the form of oil-in-water emulsion) are two of the most common adjuvants used in licensed vaccines with very well-established safety records[28, 29]. In this study, we compared Alhydrogel®and AddaVax (MF59-like adjuvant) for their ability to improve the efficacy of RBD219-N1 in mice. As shown in Figure 3 and Supplementary Table 1, mice immunized with RBD219-N1 formulated with Alhydrogel®produced potent neutralizing antibody responses, resulting in complete protection against a subsequent SARS-CoV infection. In contrast, RBD219-N1 with MF59-like adjuvant-induced high IgG titers (Supplementary Table 1) but failed to elicit protective neutralizing antibody responses and did not protect against SARS-CoV infection, as evaluated by the isolation of the infectious virus and quantitative PCR (qPCR) for viral RNA. Unlike MF59-like adjuvant formulated RBD219-N1 and RBD219-N1 alone, we also noted that RBD219-N1 formulated with Alhydrogel® did not induce a cellular infiltration within the lungs (Figure 4 and Supplementary Table 1). Taken together, these results suggest that RBD219-N1/Alhydrogel® was potentially both effective and safe, and therefore Alhydrogel® was chosen as the optimal adjuvant and further studied for the dose-ranging study.

**Table 1.** Historical lung histopathology data in mouse preclinical studies for SARS and MERS vaccines. Comparison of eosinophil infiltration induced by (A) different SARS vaccines and (B) different adjuvants for the whole virus MERS vaccine. DIV: double-inactivated whole virus vaccine, BPV: Beta propiolactone-inactivated whole virus vaccine.

| Vaccine type | Average eosinophil infiltration score | reference |
| --- | --- | --- |
| (A) SARS Vaccine |  |  |
| Study 1 (Score scale: 0-4) Score based on percent eosinophils on histologic evaluation. 7-8 mice per group. 5 microscopy fields for each mouse lung were evaluated. |  |  |
| Whole virus (DIV) | ~3 | Tseng *et al.*, 2012[12] |
| Spike protein | ~2 |  |
| Spike protein virus-like particle | >3 |  |
| Whole virus (DIV) with alum | ~2 |  |
| Spike protein with alum | ~1.5 |  |
| Spike protein virus-like particle with alum | >3 |  |
| PBS control | ~0 |  |
| Study 2 (Score scale: 0-3) Score for eosinophils as percent of infiltrating cells for each vaccine dosage group. N= 7-8 per group. 10 to 20 microscopy fields for each mouse lung were scored. Scoring: 0= ,5% of cells, 1= 5–10% of cells, 2= 10–20% of cells, 3= 20% of cells. |  |  |
| Spike protein | ~1.5 | Tseng *et al.*, 2012[12] |
| Whole virus (DIV) | ~2.5 |  |
| Whole virus (BPV) | Not available |  |
| Spike protein with alum | ~1.5 |  |
| Whole virus (DIV) with alum | ~2.5 |  |
| Whole virus (BPV) with alum | ~2.5 |  |
| PBS control | 0 |  |
| (B) MERS Vaccine |  |  |
| Study 1 (Score scale: 0-3) 0- no pathology, 1- mild, 2- moderate, and 3-severe. |  |  |
| Alum alone | 0 | Agrawal *et al.*, 2016[39] |
| MF59-like adjuvant alone | 0 |  |
| Whole virus with alum | 2 |  |
| Whole virus with MF59-like adjuvant | 2 |  |
| Whole virus | 2 |  |

**Figure 3.**
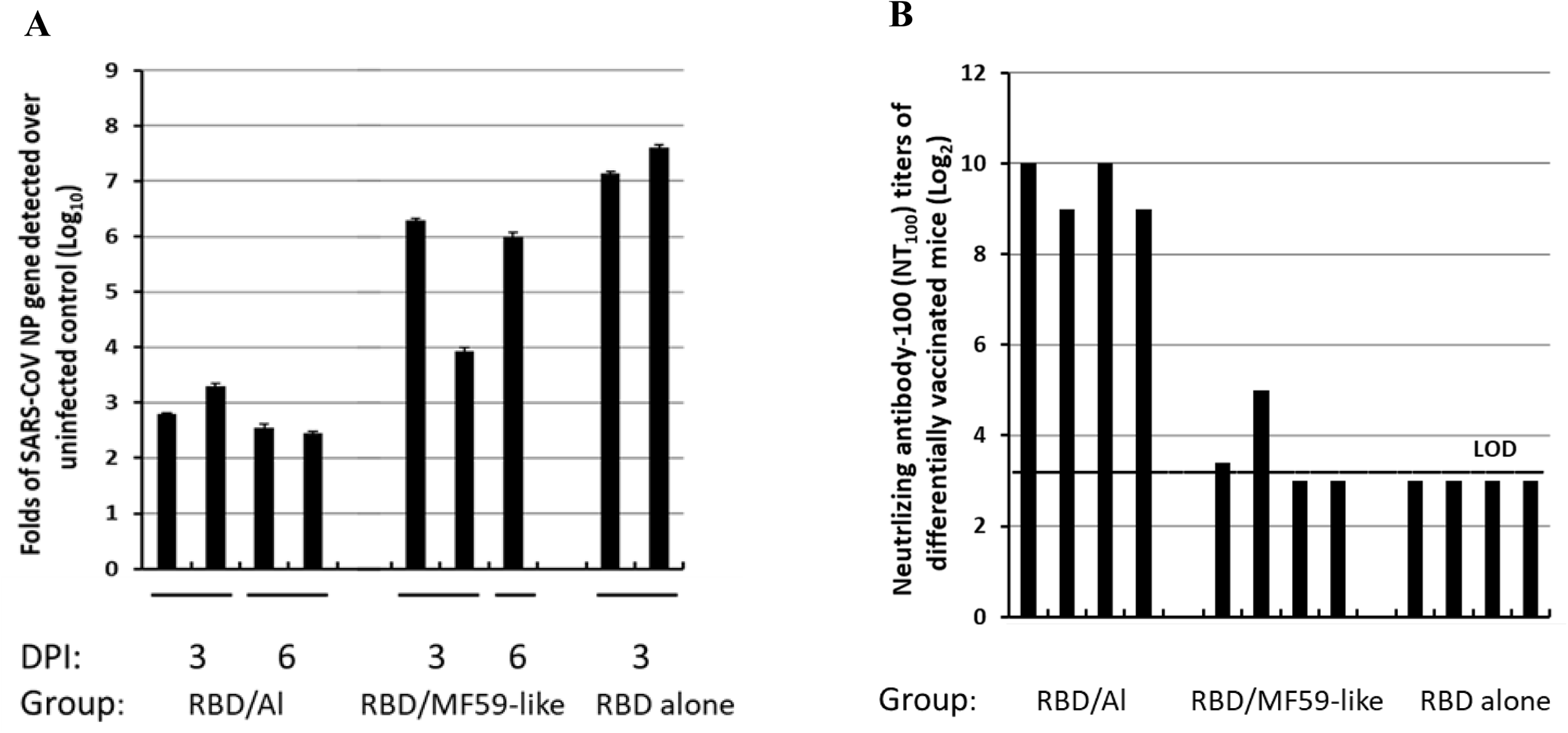
Comparison of Alhydrogel (Al) and AddaVax (MF-59-like adjuvant). (A)qPCR for the expression of the SARS-CoV np gene in the lungs of mice and (B) neutralizing antibody-100 (NT100) titers of mice differentially vaccinated with RBD219-N1 (log_2_). The mice (N=4) were vaccinated with RBD219-N1 formulated in Alhydrogel®, AddaVax (MF59-like adjuvant), and no adjuvant. On day 52, sera were collected to evaluate IgG and neutralizing antibody titers. On day 63, mice were i.n. challenged with SARS-CoV. Each bar represented an individual mice, two mice per group were sacrificed on day 3 and day 6 posted infection (dpi) to evaluate viral load.

**Figure 4.**
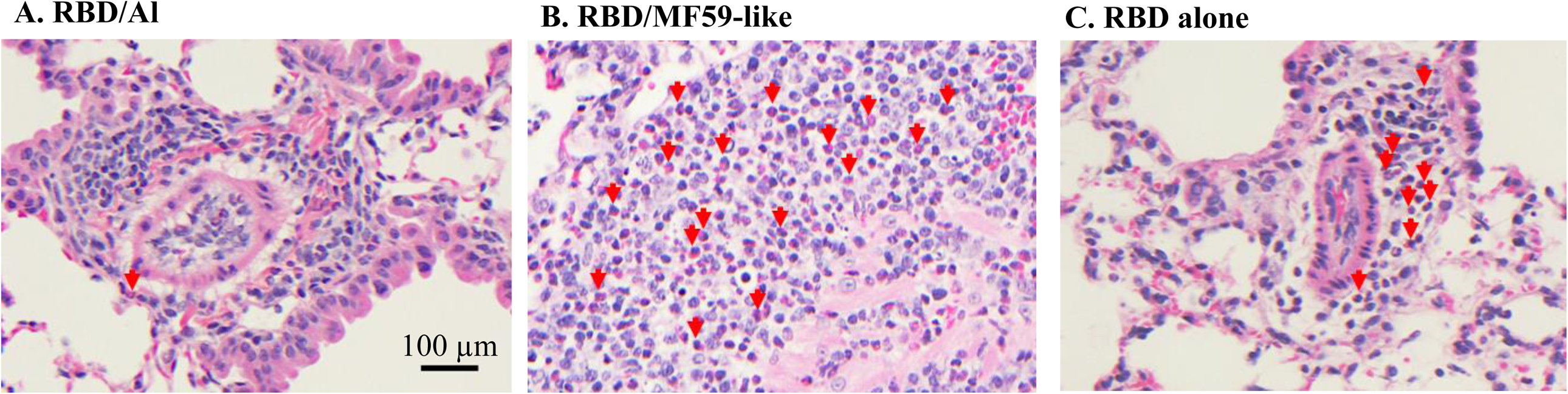
Lung histopathology of infiltrates from mice immunized with different formulations. Photomicrographs of lung tissue from Balb/c mice to evaluate eosinophil infiltration after challenge with SARS-CoV that had previously been immunized with RBD219-N1 formulated with (A) Alhydrogel® (Al), (B) AddaVax (MF59-like adjuvant), and (C) no adjuvant. (A) Alhydrogel® (Al): Perivascular mononuclear infiltrations (lymphocytes and monocytes/macrophages) along with very few eosinophils, only one seen in this field, indicated by a red arrow. (B) AddaVax (MF59-like adjuvant): Severe inflammatory infiltrations with more than 50% eosinophils, red arrows highlight some of those, and (C) no adjuvant: Inflammatory infiltrations with less than 50% eosinophils. Scale bar = 100 µm.

### Alhydrogel® dose-ranging study

Although RBD219-N1 formulated on Alhydrogel® at a ratio of 1:25 was effective in immunized animals against lethal SARS-CoV challenge without inducing apparent pulmonary immunopathology, it was of further interest to compare different aluminum ratios, including the efficacy of RBD219-N1 formulated on Alhydrogel® at both 1:25 and 1:8 ratios in mice. Sera were collected 10 days after the last vaccination to test RBD219-N1-specific IgG antibody responses by ELISA and neutralizing antibodies against live SARS-CoV infections. Results showed that mice immunized with RBD219-N1/Alhydrogel®at a 1:25 ratio produced significantly higher neutralizing antibody titers and RBD219-N1-specific IgG titers than those vaccinated at a 1:8 ratio (Figure 5A-B). Consistent with the antibody responses, mice immunized with an RBD219-N1/Alhydrogel® ratio of 1:25 were completely protected against lethal challenge with SARS-CoV, as indicated by the undetectable infectious virus within the lungs and also a lack of morbidity and mortality (Figure 6). In contrast, infectious virus was recovered from the lungs of mice immunized with a 1:8 ratio of RBD219-N1/Alhydrogel® (Groups 2 and 3) by day 5 post-challenge and one mouse from each group died. As expected, mice immunized with Alhydrogel® alone (Group 4) were not protected against a lethal viral challenge and had detectable virus in the lungs resulting in the death of 2 mice on day 4 post-challenge, while SARS-CoV S protein formulated on Alhydrogel® (Group 5) was also immunogenic and protective against SARS-CoV infection. Histopathologic examination of lung tissues using IHC (specific for eosinophils) revealed minimal eosinophilic infiltration for mice in group 1 (1:25 ratio) while somewhat increased infiltration was observed in groups 2 and 3 (1:8 ratio) and the worst eosinophils infiltration was found in the mice immunized with S protein among all groups (Supplementary Table 2 and Figure 7). The results confirm the protective efficacy of RBD219-N1/Alhydrogel®, and its superiority to the S-protein in terms of protection and reduction or prevention in eosinophilic infiltration, as well as the favorable effects of Alhydrogel®for eliciting high titer neutralizing antibody and maximal reductions in immune enhancement comparing to other groups.

**Figure 5.**
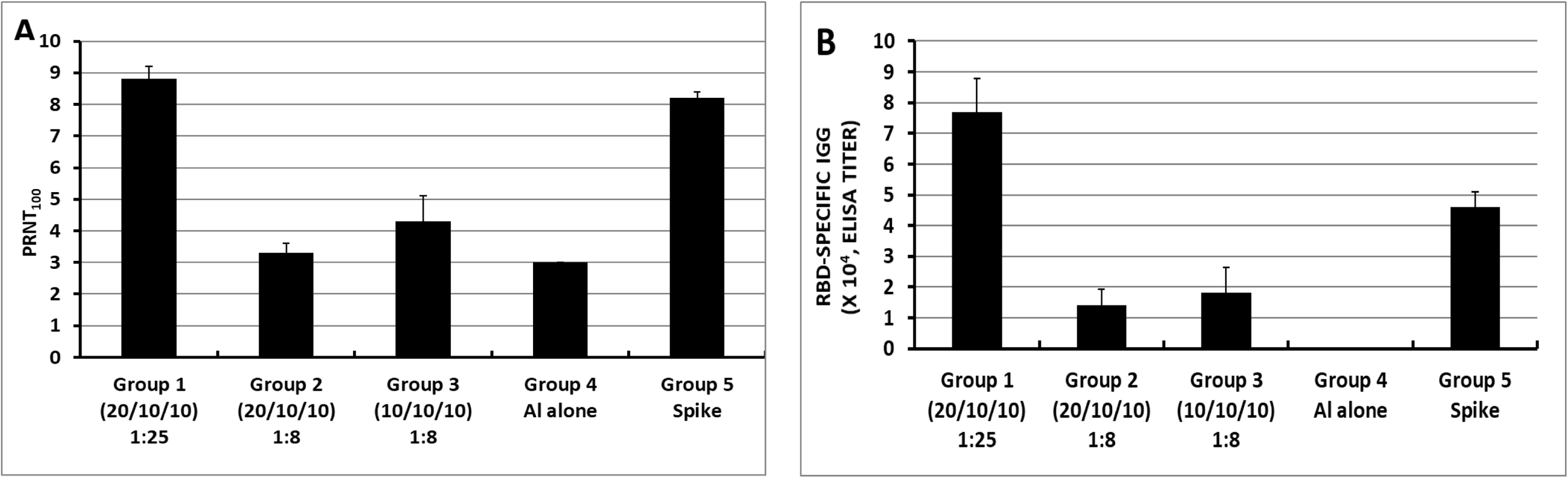
Neutralizing (A) and RBD-specific IgG antibody (B) responses of mice immunized with RBD219-N1/Alhydrogel® (Al) at 1:25 or 1:8 ratios. In group 1, a formulation of 0.2 mg/mL RBD in 5 mg/mL Alhydrogel® with the dose of 20 µg/10 µg/10 µg of RBD for prime/1st boost/2nd boost was used for immunization, respectively. In group 2, the formulation 0.2 mg/mL RBD in 1.6mg/mL Alhydrogel ®was tested, more specifically, a dosing regimen of 20 µg/10 µg/10 µg RBD for prime/1st boost/2nd boost was used while in group 3, 10 µg RBD were used for all immunizations. Alhydrogel® alone and pre-formulated SARS-S protein (3 µg) were used in groups 4 and 5, respectively, as negative and positive controls

**Figure 6.**
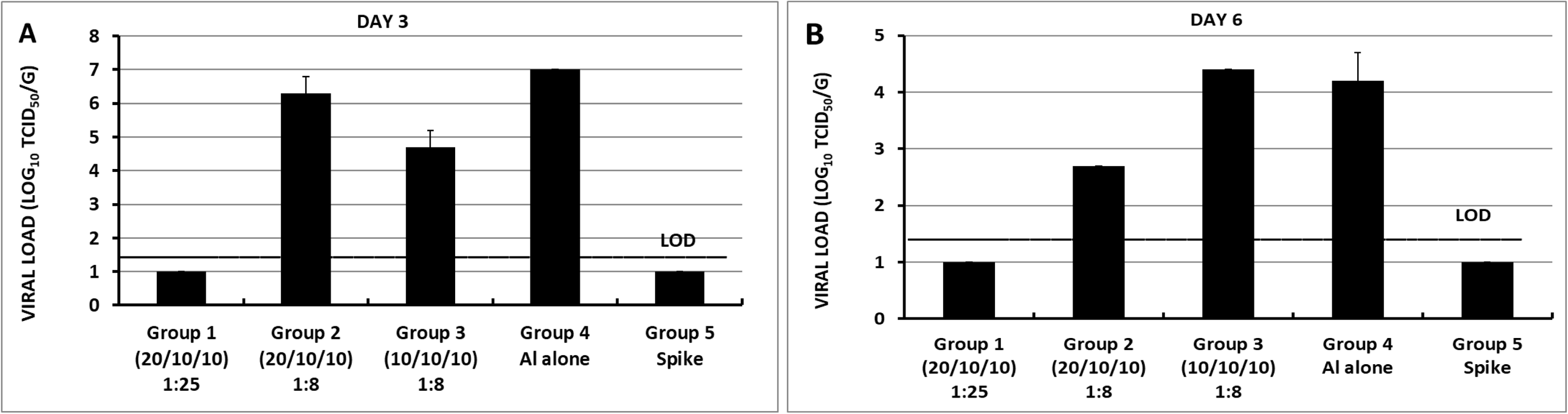
Lung viral loads of differentially immunized mice challenged intranasally with SARS-CoV on day 3 post-challenge and (B) day 6 post-challenge. In group 1, a formulation of 0.2 mg/mL RBD in 5 mg/mL Alhydrogel® with the dose of 20 µg/10 µg/10 µg of RBD for prime/1st boost/2nd boost was used for immunization, respectively. In group 2, the formulation 0.2 mg/mL RBD in 1.6mg/mL Alhydrogel ®was tested, more specifically, a dosing regimen of 20 µg/10 µg/10 µg RBD for prime/1st boost/2nd boost was used while in group 3, 10 µg RBD were used for all three immunizations. Alhydrogel® alone and pre-formulated SARS-S protein (3 µg) were used in groups 4 and 5, respectively, as negative and positive controls

**Figure 7.**
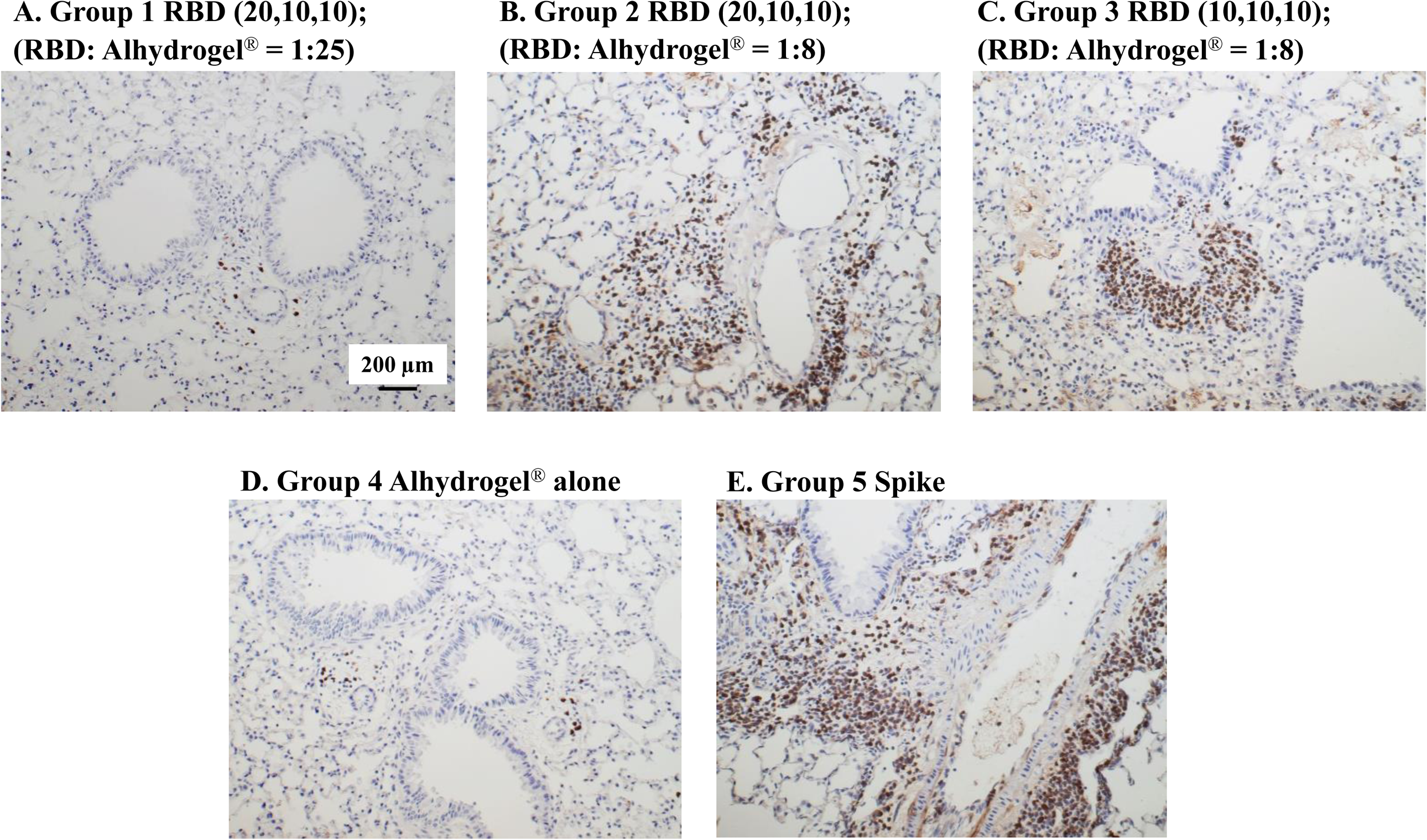
Eosinophilic infiltration in mice immunized with different doses of Alhydrogel®. Photomicrographs of lung tissue from Balb/c mice after challenge with SARS-CoV. (A) In group 1, a formulation of 0.2 mg/mL RBD in 5 mg/mL Alhydrogel®with the dose of 20 µg/10 µg/10 µg of RBD for prime/1st boost/2nd boost was used for immunization, respectively. (B) In group 2, the formulation 0.2 mg/mL RBD in 1.6mg/mL Alhydrogel® was tested with a dosing regimen of 20 µg/10 µg/10 µg RBD for prime/1st boost/2nd boost (C) In group 3, 10 µg RBD were used for all three immunizations. (D Alhydrogel® alone in group 4 was used as a negative control while (E) alum-preformulated SARS-S protein (3 µg) in group 5 was used as a positive control. Scale bar= 200 µm.

## DISCUSSION

The RBD of the S1 protein of SARS-CoV, which is responsible for the attachment of the angiotensin-converting enzyme-2 (ACE2) receptor and initiates the process for cell binding and entry, has been proven as a promising vaccine candidate [30]. It has been expressed as a recombinant protein with a hexahistidine tag, a GST protein or Fc fragment in several different expression platforms including *E. coli*, insect cells, and mammalian cells to simplify the purification process and these constructs were shown to trigger neutralizing antibodies and immunity [16, 18-21]. However, an additional tag on a recombinant protein-based vaccine should be avoided because it can potentially trigger an undesired immune response. In our previous studies [24, 25], we have expressed and purified several tag-free recombinant RBD constructs in yeast using a scalable process and discovered that the yeast-expressed RBD219-N1 induced a stronger RBD-specific antibody response and a high level of neutralizing antibodies in immunized mice when formulated with Alhydrogel®, and thus, a very promising vaccine candidate. In this study, we conducted for the first time an efficacy study in which we screened several different yeast-expressed RBD proteins and compared them to the S protein as a positive control. RBD219-N1 formulated with Alhydrogel® triggered higher neutralizing titers than the other Alhydrogel®-formulated RBD constructs or the S protein; the RBD219-N1 formulated with Alhydrogel®immunized mice were fully protected with 100% survival rate. The RBD219-N1 construct was further selected for adjuvant screening and adjuvant dose-ranging studies.

Before testing different adjuvants, we investigated the immunization routes (s.c. and i.m) for RBD219-N1 adsorbed to Alhydrogel® by evaluating the IgG antibody responses (Supplementary Figure. 1A), neutralizing antibody titer against SARS pseudovirus and live SARS-CoV infections (Supplementary Figures. 1B and 1C). It was found that both immunization routes were able to induce high titers of specific IgG and neutralizing antibodies against infections of SARS pseudovirus in ACE2/293T cells (Supplementary Figure.1B) and live SARS-CoV in Vero cells (Supplementary Figure.1C). Although the antibody responses induced through the s.c. route were significantly higher than those through the i.m. route, the i.m. route was selected for subsequent adjuvant optimization because i.m. injection of the vaccines containing adjuvants has less chance to induce adverse local effects than s.c. injection[31] and the majority of the clinically used vaccines are administered via the i.m. route [32].

The superiority of the RBD219-N1 vaccine antigen to the S protein was reflected both in terms of eliciting neutralizing antibodies and protective immunity, and that the alum-adjuvanted RBD219-N1 resulted in little to no cellular or eosinophilic immunopathology compared to either the S-protein or an M59-like adjuvant-formulated RBD219-N1 vaccine. With regards to the former observation, it was previously shown that removal of immune-enhancing epitopes located outside the S-protein RBD domain may result in an immunogen, which is less likely to induce immunopathology [15, 33]. This finding led to the initial selection of the RBD as a vaccine antigen [34]. With regards to adjuvant selection, alum prevented or reduced immune enhancement, a finding that confirms a previous observation for the doubly inactivated virus, viral-like particle vaccines, and S protein[12]. For example, Tseng et al. reported that mice immunized with aluminum-formulated virus-like-particle and inactivated virus vaccines showed less immunopathology than the non-adjuvanted vaccines. Such findings provided the basis for suggesting that eosinophilic immunopathology may occur through mechanisms other than Th2 immunity [35, 36].

To understand the mechanisms of potential immunopathology linked to SARS vaccines, it’s been shown that high levels of eosinophilic immunopathology were observed with modified vaccinia virus Ankara-based vaccine platform vaccines,[6-11] and this vaccine platform was found to induce both Th1 (IFN-gamma, IL-2) and Th2 (IL-4, IL-5) cytokines and down-regulation of anti-inflammatory cytokines (IL-10, TGF-beta) upon infection, causing severe infiltration of neutrophils, eosinophils, and lymphocytes into the lung. These pieces of evidence suggest that Th2 is not the sole factor but rather a mixed Th1 and Th2 response is responsible for the immunopathology. Additionally, it was found that IL-6 was shown to have a prominent role in SARS-CoV-induced immune enhancement [11] in experimental animals, as well as in lung pathology in SARS patients [37]. The prominent role of IL-6 in host Th17 immune responses suggest that this pathway might also comprise a component of coronavirus vaccine immune enhancement[13]. The finding that Th17 lymphocytes activate eosinophils [35], and that eosinophils comprise a key element of Th17 responses is consistent with these findings [35]. Of interest, monoclonal antibodies directed at interfering with IL-6 binding with its receptor are now being investigated as possible immunotherapies for patients with COVID19 [38].

Finally, in the Alhydrogel®dose-ranging study, we observed that a higher concentration of Alhydrogel® was required to trigger a fully protective immune response with attenuated eosinophils infiltration, while S protein induced a higher degree of eosinophilic cellular infiltration, which is consistent with the previous finding [12]. Table 1A further summarized the comparison of eosinophilic immunopathology induced by different SARS-CoV vaccines, including the S protein, virus-like particle (VLP) expressing spike protein, and the inactivated whole virus vaccines. The results indicated that the vaccine using VLP expressing S protein triggered worse eosinophilic infiltration than the inactivated whole virus vaccines, and S protein showed the least eosinophil infiltration among them either with or without formulated with alum. Our studies find that the RBD can elicit far less immunopathology than even the S protein. Similar to what was seen for the whole virus vaccine against SARS, lung eosinophil infiltration was observed in mice immunized with MERS gamma irradiation-inactivated whole virus vaccine after challenge with MERS-CoV (Table 1B) [39]. These findings further suggested that Alhydrogel®formulated RBD219-N1 was a safer SARS-CoV vaccine than the ones listed in Table 1A.

Collectively, all the preclinical data suggested that SARS-CoV RBD219-N1 formulated with alum is a potentially safe and efficacious vaccine against SARS-CoV infection. We have developed the scalable process and partnered with WRAIR and manufactured RBD219-N1 protein under current good manufacturing practices (cGMP) in 2016. The bulk drug substance has been frozen (−70°C to -80°C) in a temperature-regulated storage location and under stability testing since its manufacturing date (July 2016) and remains stable. This vaccine candidate is ready for formulation and can be rapidly transitioned to clinical testing. The preclinical data reported here suggest that SARS-CoV RBD219-N1 formulated with Alhydrogel®is a safer and more efficacious vaccine against SARS-CoV infection compared to many other candidate vaccines. This vaccine is also under evaluation as part of a broader strategy to accelerate as a universal CoV vaccine, possibly in combination with RBDs from other coronaviruses, including SARS-CoV-2.

## ACKNOWLEDGMENT

This study was supported by a grant from the National Institutes of Health (R01AI098775).

## SUPPLEMENTAL DATA

**Supplementary Table 1**. Immunogenicity and efficacy results for adjuvant screening. n/s: not significant, M: moderate, S: severe. ND: Not detected. N/A: not available

**Supplementary Table 2**. Immunogenicity and efficacy results for the Alhydrogel dose-ranging study.

* Sacrifice; ** Not applicable; # Not detected; § Death due to over anesthetization

**Supplementary Figure 1. Optimization of immunization routes**. Mice were immunized with RBD219-N1 formulated with or without Alhydrogel® (1:25 ratio) subcutaneously (s.c.) or intramuscularly (i.m.), three times, at 3-week intervals. Sera were collected 10 days after the last immunization and tested for IgG antibody responses and for neutralizing antibodies against SARS pseudovirus and live SARS-CoV infections. (A) Detection of IgG antibody response by ELISA in mouse sera. Neutralization antibody titers against SARS pseudovirus

(B)and live SARS-CoV (C) in mouse sera. [Alhydrogel® abbreviated as Alum.]. Psudovirus was prepared as previously described in Chen et al., 2014 [20].

## REFERENCES

[1] Zhong NS, Zheng BJ, Li YM, Poon, Xie ZH, Chan KH, et al. Epidemiology and cause of severe acute respiratory syndrome (SARS) in Guangdong, People’s Republic of China, in February, 2003. Lancet. 2003;362:1353–8.

[2] Drosten C, Gunther S, Preiser W, van der Werf S, Brodt HR, Becker S, et al. Identification of a novel coronavirus in patients with severe acute respiratory syndrome. N Engl J Med. 2003;348:1967–76.

[3] Ksiazek TG, Erdman D, Goldsmith CS, Zaki SR, Peret T, Emery S, et al. A novel coronavirus associated with severe acute respiratory syndrome. N Engl J Med. 2003;348:1953–66.

[4] Zaki AM, van Boheemen S, Bestebroer TM, Osterhaus AD, Fouchier RA. Isolation of a novel coronavirus from a man with pneumonia in Saudi Arabia. N Engl J Med. 2012;367:1814–20.

[5] Du L, He Y, Zhou Y, Liu S, Zheng B-J, Jiang S. The spike protein of SARS-CoV--a target for vaccine and therapeutic development. Nature reviews Microbiology. 2009;7:226–36.

[6] Perlman S, Dandekar AA. Immunopathogenesis of coronavirus infections: implications for SARS. Nat Rev Immunol. 2005;5:917–27.

[7] Bolles M, Deming D, Long K, Agnihothram S, Whitmore A, Ferris M, et al. A double-inactivated severe acute respiratory syndrome coronavirus vaccine provides incomplete protection in mice and induces increased eosinophilic proinflammatory pulmonary response upon challenge. J Virol. 2011;85:12201–15.

[8] Weingartl H, Czub M, Czub S, Neufeld J, Marszal P, Gren J, et al. Immunization with modified vaccinia virus Ankara-based recombinant vaccine against severe acute respiratory syndrome is associated with enhanced hepatitis in ferrets. J Virol. 2004;78:12672–6.

[9] Czub M, Weingartl H, Czub S, He R, Cao J. Evaluation of modified vaccinia virus Ankara based recombinant SARS vaccine in ferrets. Vaccine. 2005;23:2273–9.

[10] Liu L, Wei Q, Lin Q, Fang J, Wang H, Kwok H, et al. Anti-spike IgG causes severe acute lung injury by skewing macrophage responses during acute SARS-CoV infection. JCI Insight. 2019;4.

[11] Yasui F, Kai C, Kitabatake M, Inoue S, Yoneda M, Yokochi S, et al. Prior immunization with severe acute respiratory syndrome (SARS)-associated coronavirus (SARS-CoV) nucleocapsid protein causes severe pneumonia in mice infected with SARS-CoV. J Immunol. 2008;181:6337–48.

[12] Tseng CT, Sbrana E, Iwata-Yoshikawa N, Newman PC, Garron T, Atmar RL, et al. Immunization with SARS coronavirus vaccines leads to pulmonary immunopathology on challenge with the SARS virus. PLoS One. 2012;7:e35421.

[13] Hotez PJ, Bottazzi ME, Corry DB. The Potential Role of Th17 Immune Responses in Coronavirus Immunopathology and Vaccine-induced Immune Enhancement. Microbes and Infection. 2020.

[14] Jaume M, Yip MS, Kam YW, Cheung CY, Kien F, Roberts A, et al. SARS CoV subunit vaccine: antibody-mediated neutralisation and enhancement. Hong Kong Med J. 2012;18 Suppl 2:31–6.

[15] Wang Q, Zhang L, Kuwahara K, Li L, Liu Z, Li T, et al. Immunodominant SARS Coronavirus Epitopes in Humans Elicited both Enhancing and Neutralizing Effects on Infection in Non-human Primates. ACS Infect Dis. 2016;2:361–76.

[16] Wong SK, Li W, Moore MJ, Choe H, Farzan M. A 193-amino acid fragment of the SARS coronavirus S protein efficiently binds angiotensin-converting enzyme 2. J Biol Chem. 2004;279:3197–201.

[17] Jiang S, Bottazzi ME D. L, Lustigman S, Tseng C-TK, Curti E, et al. Roadmap to developing a recombinant coronavirus S protein receptor-binding domain vaccine for severe acute respiratory syndrome. Expert Review of Vaccines. 2012;11:1405–13.

[18] Du L, Zhao G, Chan CC, Sun S, Chen M, Liu Z, et al. Recombinant receptor-binding domain of SARS-CoV spike protein expressed in mammalian, insect and E. coli cells elicits potent neutralizing antibody and protective immunity. Virology. 2009;393:144–50.

[19] Du L, Zhao G, Chan CC, Li L, He Y, Zhou Y, et al. A 219-mer CHO-expressing receptor-binding domain of SARS-CoV S protein induces potent immune responses and protective immunity. Viral Immunol. 2010;23:211–9.

[20] Du L, Zhao G, Li L, He Y, Zhou Y, Zheng BJ, et al. Antigenicity and immunogenicity of SARS-CoV S protein receptor-binding domain stably expressed in CHO cells. Biochem Biophys Res Commun. 2009;384:486–90.

[21] Du L, Zhao G, He Y, Guo Y, Zheng BJ, Jiang S, et al. Receptor-binding domain of SARS-CoV spike protein induces long-term protective immunity in an animal model. Vaccine. 2007;25.

[22] He Y, Lu H, Siddiqui P, Zhou Y, Jiang S. Receptor-binding domain of severe acute respiratory syndrome coronavirus spike protein contains multiple conformation-dependent epitopes that induce highly potent neutralizing antibodies. J Immunol. 2005;174:4908–15.

[23] He Y, Li J, Li W, Lustigman S, Farzan M, Jiang S. Cross-neutralization of human and palm civet severe acute respiratory syndrome coronaviruses by antibodies targeting the receptor-binding domain of spike protein. J Immunol. 2006;176:6085–92.

[24] Chen WH, D. L, Chag SM, Ma C, Tricoche N, Tao X, et al. Yeast-expressed recombinant protein of the receptor-binding domain in SARS-CoV spike protein with deglycosylated forms as a SARS vaccine candidate. Hum Vaccin Immunother. 2014;10:648–58.

[25] Chen WH, Chag SM, Poongavanam MV, Biter AB, Ewere EA, Rezende W, et al. Optimization of the Production Process and Characterization of the Yeast-Expressed SARS-CoV Recombinant Receptor-Binding Domain (RBD219-N1), a SARS Vaccine Candidate. J Pharm Sci. 2017;106:1961–70.

[26] Roberts A, Deming D, Paddock CD, Cheng A, Yount B, Vogel L, et al. A mouse-adapted SARS-coronavirus causes disease and mortality in BALB/c mice. PLoS pathogens. 2007;3:e5.

[27] Tseng CT, Huang C, Newman P, Wang N, Narayanan K, Watts DM, et al. Severe acute respiratory syndrome coronavirus infection of mice transgenic for the human Angiotensin-converting enzyme 2 virus receptor. J Virol. 2007;81:1162–73.

[28] Kommareddy S, Singh M, O’Hagan DT. Chapter 13 - MF59: A Safe and Potent Adjuvant for Human Use. In: Schijns VEJC, O’Hagan DT, editors. Immunopotentiators in Modern Vaccines (Second Edition): Academic Press; 2017. p. 249–63.

[29] Garçon N, Friede M. 6 - Evolution of Adjuvants Across the Centuries. In: Plotkin SA, Orenstein WA, Offit PA, Edwards KM, editors. Plotkin’s Vaccines (Seventh Edition): Elsevier; 2018. p. 61-74.e4.

[30] Jiang S, Bottazzi ME, D. L, Lustigman S, Tseng CT, Curti E, et al. Roadmap to developing a recombinant coronavirus S protein receptor-binding domain vaccine for severe acute respiratory syndrome. Expert Rev Vaccines. 2012;11:1405–13.

[31] Zuckerman JN. The importance of injecting vaccines into muscle. Different patients need different needle sizes. BMJ. 2000;321:1237–8.

[32] Center_for_Disease_Control_and_Prevention. Administer the vaccine(s).

[33] Jiang S, He Y, Liu S. SARS vaccine development. Emerg Infect Dis. 2005;11:1016–20.

[34] Du L, Zhao G, He Y, Guo Y, Zheng BJ, Jiang S, et al. Receptor-binding domain of SARS-CoV spike protein induces long-term protective immunity in an animal model. Vaccine. 2007;25:2832–8.

[35] Dias PM, Banerjee G. The role of Th17/IL-17 on eosinophilic inflammation. J Autoimmun. 2013;40:9–20.

[36] Keely S, Foster PS. Stop Press: Eosinophils Drafted to Join the Th17 Team. Immunity. 2015;43:7–9.

[37] Zhang Y, Li J, Zhan Y, Wu L, Yu X, Zhang W, et al. Analysis of serum cytokines in patients with severe acute respiratory syndrome. Infect Immun. 2004;72:4410–5.

[38] Clinicaltrials.gov. Tocilizumab in COVID-19 Pneumonia (TOCIVID-19) (TOCIVID-19). (2000, Mar. 19-). 2020.

[39] Agrawal AS, Tao X, Algaissi A, Garron T, Narayanan K, Peng BH, et al. Immunization with inactivated Middle East Respiratory Syndrome coronavirus vaccine leads to lung immunopathology on challenge with live virus. Hum Vaccin Immunother. 2016;12:2351–6.

