## Supplementary material for "Yeast-Expressed SARS-CoV Recombinant Receptor-Binding Domain (RBD219-N1) Formulated with Aluminum Hydroxide Induces Protective Immunity and Reduces Immune Enhancement": sup. data

**SUPPLEMENTARY DATA**

**Supplementary Table 1.** Immunogenicity and efficacy results for adjuvant screening. n/s: not significant, M: moderate, S: severe. ND: Not detected. N/A: not available

| Vaccination Groups | $NT_{100}$ ($log_2$) | $NT_{50}$ ($Log_2$) | infectious virus/gram (dpi) | Infiltration grades (dpi) | EOS# (dpi) | Specific IgG titers (x $10^3$) (ELISA) | IgG1/IgG2a ratios |
|---|---|---|---|---|---|---|---|
| RBD219-N1/Alhydrogel | 1,280 (10) | --* | ND** (3) | 1 (3) | n/s (3) | 512 | 4 |
| | 640 (9) | 1,280 (10) | ND (3) | | | 512 | 8 |
| | 1,280 (10) | 2,560 (11) | ND (6) | 1 (6) | n/s (6) | 128 | 4 |
| | 640 (9) | 1,280 (10) | ND (6) | | | 256 | 8 |
| RBD219-N1/MF59-like adjuvant | N/A | 20 | $7.5 \times 10^4$ (3) | 3 (3) | S | 128 | 64 |
| | 80 | -- | $5 \times 10^4$ (3) | | | 256 | 1,024 |
| | N/A | < 20 | ND (6) | 3 (6) | S | 512 | 64 |
| | N/A | < 20 | Dead (6) | | | 512 | 256 |
| RBD219-N1 alone | N/A | < 20 | $5 \times 10^6$ (3) | 2 (3) | M | 0.1 | -- |
| | N/A | < 20 | $7.5 \times 10^6$ (3) | | | ND | -- |
| | N/A | < 20 | Dead (4) | -- | -- | 0.4 | -- |
| | N/A | < 20 | Dead (5) | | | ND | -- |

**Supplementary Table 2.** Immunogenicity and efficacy results for Alhydrogel dose-ranging study.
* Sacrifice; ** Not applicable; # Not detected; § Death due to over anesthetization

| Groups | Animal ID# | $NT_{100}$ | $NT_{50}$ | ELISA titers RBD-specific IgG antibody | RBD-specific IgG1/IgG2a antibody ratio | Infectious virus/g (dpi) | Eos-infiltration grade |
|---|---|---|---|---|---|---|---|
| RBD219-N1/ Alhydrogel (RBD: 20, 10, 10 µg; Alhydrogel: 500, 250, 250 µg) | 1 | > 1,280 | NA** | 102,400 | 32 | ND# (3) | 1 |
| | 2 | 640 | NA | 102,400 | 16 | Death (OA)§ | NA |
| | 3 | 640 | 1,280 | 102,400 | 16 | ND (3) | 1 |
| | 4 | 160 | 320 | 51,200 | 8 | ND (6) | 1 |
| | 5 | 320 | 640 | 51,200 | 16 | ND (6) | 1 |
| | 6 | > 1,280 | NA | 51,200 | 32 | Death (OA) | NA |
| RBD219-N1/ Alhydrogel (RBD: 20, 10, 10 µg; Alhydrogel: 160, 80, 80 µg) | 7 | < 10 | NA | 6,400 | 64 | $1 \times 10^6$ (3) | 2 |
| | 8 | < 10 | NA | 1,600 | 32 | $2.5 \times 10^6$ (3) | 2 |
| | 9 | < 10 | NA | 200 | NA | $2.5 \times 10^6$ (3) | 2 |
| | 10 | < 10 | NA | 25,600 | 256 | Dead (6) | NA |
| | 11 | < 10 | NA | 25,600 | 64 | $5 \times 10^4$ (6) | 2 |
| | 12 | 40 | 80 | 25,600 | 64 | ND (6) | 2 |
| RBD219-N1/ Alhydrogel (RBD: 10, 10, 10 µg; Alhydrogel: 80, 80, 80 µg) | 13 | < 10 | NA | 6,400 | 128 | $1 \times 10^5$ (3) | 2 |
| | 14 | < 10 | NA | 400 | NA | Dead (3) | NA |
| | 15 | 160 | 320 | 25,600 | 64 | $1 \times 10^4$ (3) | 2 |
| | 16 | 160 | NA | 51,200 | 64 | ND (6) | 2 |
| | 17 | 10 | 20 | 25,600 | 4 | ND (6) | 2 |
| | 18 | < 10 | NA | 800 | 32 | $2.5 \times 10^4$ (6) | 2 |
| TRIS-Alhydrogel (Alhydrogel: 160, 160, 160 µg) | 19 | < 10 | NA | < 100 | NA | Dead (3) | NA |
| | 20 | < 10 | NA | < 100 | NA | Dead (3) | NA |
| | 21 | < 10 | NA | < 100 | NA | $1 \times 10^7$ (3) | 1 |
| | 22 | < 10 | NA | < 100 | NA | $5 \times 10^5$ (3) | 1 |
| | 23 | < 10 | NA | < 100 | NA | $2.5 \times 10^4$ (6) | 1 |
| | 24 | < 10 | NA | < 100 | NA | $1 \times 10^4$ (6) | 1 |
| SARS-S/Alum (S: 3, 3, 3 µg; Alum: pre-formulated) | 25 | 640 | NA | 51,200 | 32 | ND (3) | 2+ |
| | 26 | 320 | NA | 25,600 | 16 | ND (3) | 2+ |
| | 27 | 320 | NA | 51,200 | 32 | ND (3) | 2+ |
| | 28 | 320 | NA | 51,200 | 32 | ND (6) | 2+ |
| | 29 | 320 | NA | 51,200 | 32 | ND (6) | 2+ |
| | 30 | Sac* | NA | NA | NA | NA | NA |

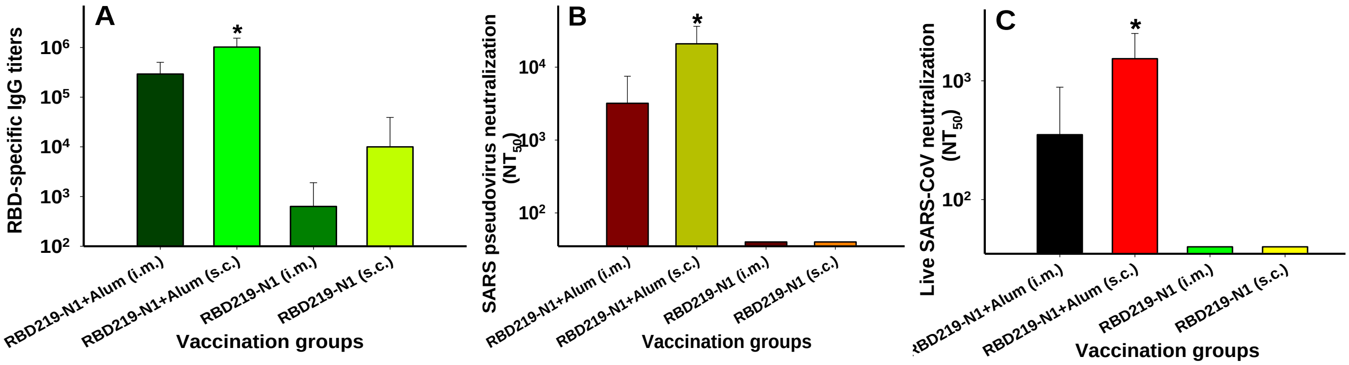


**Supplementary Figure 1. Optimization of immunization routes.** Mice were immunized with RBD219-N1 formulated with or without Alhydrogel® (1:25 ratio) subcutaneously (s.c.) or intramuscularly (i.m.), three times, at 3-week intervals. Sera were collected 10 days after the last immunization and tested for IgG antibody responses and for neutralizing antibodies against SARS pseudovirus and live SARS-CoV infections. (A) Detection of IgG antibody response by ELISA in mouse sera. Neutralization antibody titers against SARS pseudovirus (B) and live SARS-CoV (C) in mouse sera. [Alhydrogel® abbreviated as Alum.]. Psudovirus was prepared as previously described in Chen et al., 2014 [20].
